# Dopamine projections to the basolateral amygdala enable reward prediction

**DOI:** 10.64898/2026.06.21.733629

**Authors:** Kathia Ramírez-Armenta, Christina Feng, Andrea Martinez, Jalen E. Andrade, Jasmine Quach, Natalie Paredes, Ana C. Sias, Nicholas K. Griffin, Melissa J. Sharpe, Kate M. Wassum

**Author notes:** Correspondence: Kate Wassum, Dept. of Psychology, UCLA, 1285 Franz Hall, Box 951563, Los Angeles, CA 90095-1563.

## Abstract

Reward predictions are critical to both adaptive learning and decision making. Such predictions are supported by environmental cues that signal the availability and identity of rewarding events. Here we used fiber photometry, cell-type and pathway-specific optogenetic inhibition, Pavlovian cue–reward conditioning, and decision-making tests in male and female rats to reveal that ventral tegmental area dopamine (VTA_DA_) projections to the basolateral amygdala (BLA) support cue–reward predictions. Reward-predictive cues trigger dopamine release in the BLA that encodes the value of the predicted reward. This cue-evoked VTA_DA_→BLA activity mediates the ability of cue-reward predictions to bias action selection and adapt cue-response decisions based on the current value of the predicted reward. Cue-evoked VTA_DA_→BLA activity also mediates the constraining influence of cue-reward predictions on new learning. Thus, cue-evoked BLA dopamine supports the reward predictions that both enable adaptive decision making and constrain learning.

---

Our brains are prediction machines. They use prior experience to build an internal model of environmental relationships that enables us to predict the identity and value of forthcoming events to ensure adaptive learning and decision making^1, 2^. Cue-reward associations are fundamental components of these models^3, 4^. Whether homeostatic, social, economic, or another form of reward-related decision making, cue-reward associations allow us to use predictive environmental cues to generate the predictions and inferences needed for flexible, advantageous choices^3–6^. For example, restaurant logos at your local food court signal the availability of specific types of food (e.g. pizza, salad), allowing you to consider your options and choose the most optimal lunch in your current state (e.g., do you need something light or filling? did you become ill the last time you ate salad?). Predictions also constrain learning. When the brain is able to fully predict a forthcoming event, it will be less likely to learn a new, redundant predictor of that event^7–9^. If you already know a particular logo signals pizza availability, you will be unlikely to learn the new employee uniform also predicts pizza. Thus, predictions are important to both how we learn and how we decide. But much is unknown of the brain systems that support reward prediction.

Dopamine has long been known to be vital for learning. Midbrain dopamine neurons respond to unexpected rewards and can signal errors in reward prediction^10–13^. This signal supports predictive learning^12, 14, 15^, including recent evidence of identity-specific cue-reward learning^16–18^. In this view, dopamine is primarily a teaching signal, rather than an online contributor to reward prediction. But there are also long-standing views that dopamine functions online to support motivated behavior^19–22^. Reward-predictive cues activate dopamine neurons^23–25^ and this scales with predicted reward value and probability^26–28^. This activity can support higher-order learning^15^, but is also positioned to modulate the brain’s ability to use those cues to generate predictions. Dopamine may support cue-reward prediction via its release in the basolateral amygdala (BLA). The BLA is activated by reward cues^29–36^ and this supports the influence of cue-reward predictions on adaptive decision making^36–42^. The BLA receives input from ventral tegmental area dopamine (VTA_DA_) neurons ^43–45^ and, with learning, reward cues can evoke dopamine release in BLA^16, 46^. Therefore, here we asked whether VTA_DA_→BLA neurons supports reward prediction by combining BLA dopamine recording and inhibition with behavioral tests of the influence of identity-specific cue-reward predictions over both decision making and learning.

## RESULTS

### Cue-evoked basolateral amygdala dopamine release encodes predicted reward value during decision making

#### Cues trigger dopamine release in the BLA during decision making

We first monitored BLA dopamine release as cues influenced decision making (Figure 1a-b). Male and female rats were food restricted and received instrumental conditioning in which each of two different lever-press actions earned one of two distinct food rewards (e.g., left press→ensure/right press→pellets; 11 sessions; Figure 1c). All rats acquired the instrumental lever-press behavior (Figure 1d). To engender cue-reward associations, rats then received Pavlovian conditioning during which 2 distinct, 30-s auditory cues (aka, conditioned stimuli) each predicted delivery of one of the unique food rewards at cue offset (e.g., white noise—ensure/click—pellets; Figure 1c). With training, all rats developed a Pavlovian conditional goal-approach response (Figure 1e). To evaluate the influence of the cue-reward predictions on decision making, we next gave subjects an outcome-specific Pavlovian-to-instrumental transfer (PIT) test (Figure 1f). During this test both levers were present, but pressing was not reinforced. Each cue was presented 4 times (also without accompanying reward), with intervening cue-free baseline periods. Because the cues are never directly associated with the instrumental actions, this test assesses the ability to use the cues to generate reward predictions that influence action selection and performance in a novel choice scenario^47, 48^. If subjects are using the cues to predict specific forthcoming rewards, then during cue presentation they should preferentially press on the lever that, during training, earned the same reward as predicted by that cue, relative to the lever associated with the different reward. Indeed, rats showed this outcome-specific PIT effect (Figure 1g-h), in addition to conditional food-port approach responses (Figure 1i).

**Figure 1.**
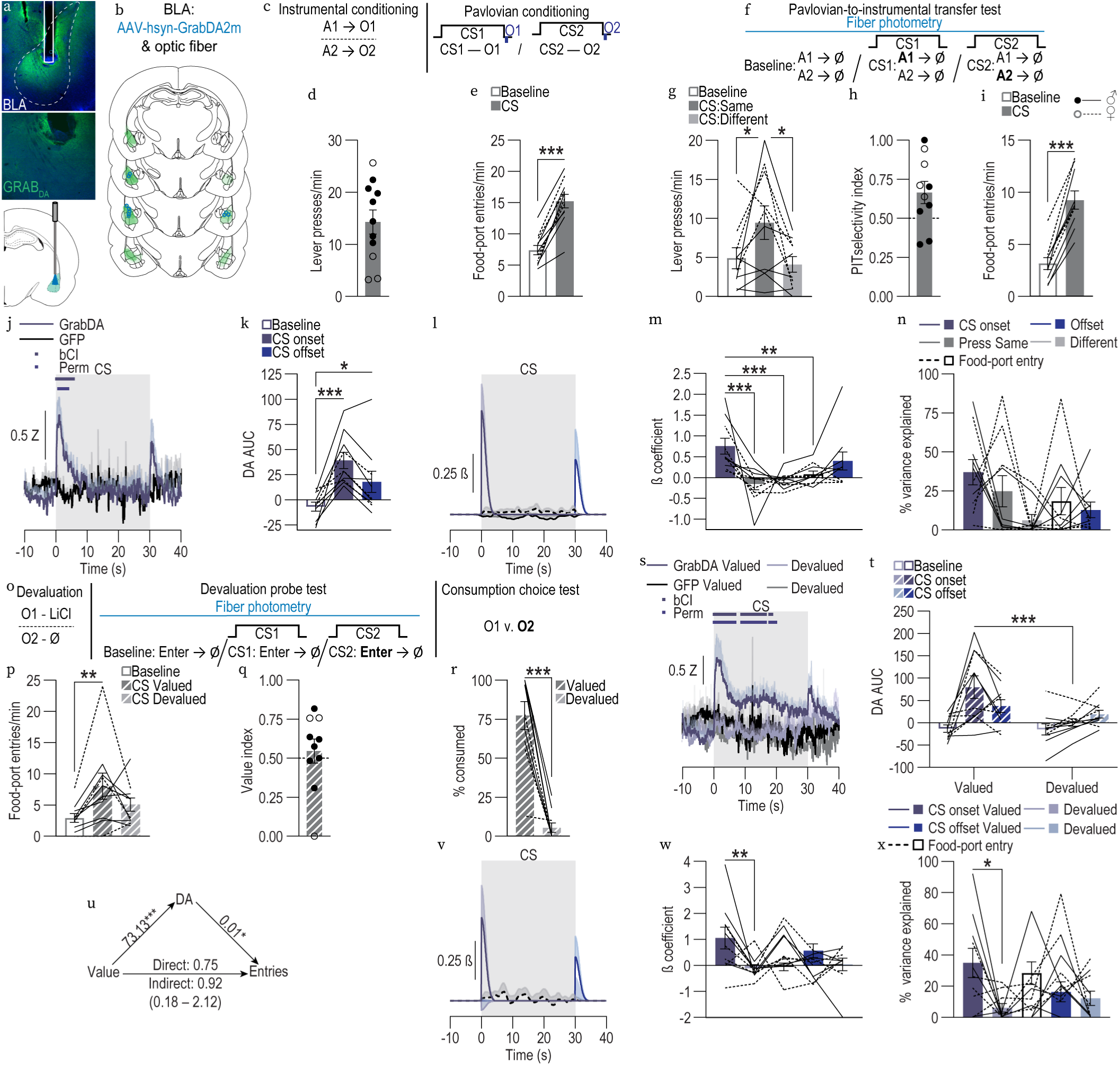
Cue-triggered basolateral amygdala dopamine release encodes predicted reward value during decision making. **(a)** Bottom: Fiber photometry approach for imaging GRABDA fluorescence changes in BLA neurons. Top: Representative fluorescent image of BLA GRABDA2m expression and fiber placement. **(b)** Schematic representation of GRABDA2m expression and optical fiber tips in BLA for all subjects. **(c-e)** Instrumental and Pavlovian conditioning **(c)** Conditioning procedures. A, action (left or right lever press); CS, 30-s conditioned stimulus (aka, “cue”, white noise or click) followed immediately by reward outcome (O, ensure solution or grain pellet). **(d)** Lever-press rate averaged across levers and across the final 2 instrumental sessions. **(e)** Food-port entry rate during averaged across cues and across the final 2 Pavlovian conditioning sessions. 2-tailed students t-test, t10 = 10.58, *P* < 0.0001, 95% CI 6.19 - 9.50. **(f-n)** Outcome-specific Pavlovian-to-instrumental transfer test. **(f)** PIT procedure. Ø, no reward was delivered. **(g)** Lever-press rates on the lever that earned the “Same” outcome as predicted or current cue or on the other available lever (Different) compared to pre-cue baseline press rate averaged across levers. 1-way repeated measures ANOVA, CS/Lever: F2,18 = 5.31, *P* = 0.02. **(h)** PIT selectivity index [Same presses/(Same + Different presses)]. 2-tailed students t-test against hypothetical no change value of 0.5, t9 = 2.33, *P* = 0.04, 95% CI - 0.33 to −0.005. **(i)** Food-port entry rate. 2-tailed students t-test, t9 = 10.24, *P* < 0.0001, 95% CI 4.75 - 7.44. **(j)** GRABDA and control GFP fluorescence changes (Z-score) during the cues during the PIT test. Waveform analysis: bCI, time points where the adjusted 95% bootstrap confidence interval for the difference between GRABDA and GFP excludes zero (10-sample cluster minimum); Perm, time points for which GRABDA is significantly (*P* < 0.05) different from GFP from permutation test. **(k)** Area under the BLA GRABDA Z-scored curve (AUC). 1-way repeated measures ANOVA, CS period: F2,18 = 14.85, *P* = 0.0002. **(l-n)** Linear regression model using convolved CS onset, offset, Same and Different lever presses, and food-port entries to predict GRABDA signal at each time point 10 s prior to and 40 s after each cue onset from the event kernels. Average regression coefficient of each kernel around cue presentation (l), β coefficients from model (m; 1-way repeated measures ANOVA, CS period: F4,36 = 6.94, *P* = 0.0003), and percentage of variance explained by each event kernel (n; 1-way repeated measures ANOVA, CS period: F4,36 = 1.98, *P* = 0.12). PIT, GRABDA: *N* = 10 (6 male); GFP: *N* = 4 (2 male). **(o-x)** Outcome-specific devaluation test. **(o)** Devaluation procedures. LiCl, lithium chloride 0.3M, 1.5% volume/weight. **(p)** Food-port entry rates during the 35-s pre-cue baseline periods compared to that during the cue (30-s cue periods + 5-s post-cue periods) predicting the non-devalued (Valued) and Devalued outcome. 1-way repeated measures ANOVA, CS/Value: F2,18 = 5.48, *P* = 0.01. **(q)** Value index [Valued CS entries/(Valued + Devalued CS entries)]. 2-tailed students t-test against hypothetical no change value of 0.5, t9 = 0.59, *P* = 0.57, 95% CI −0.22 - 0.13. **(r)** Post-devaluation consumption test. % of total available food (10 g or 10 mL) consumed during 10-min test. 2-tailed students t-test, t9 = 7.87, *P* < 0.0001, 95% CI −92.52 - −51.20. **(s)** GRABDA and control GFP fluorescence changes (Z-score) during the cues during the devaluation probe test. Waveform analysis: bCI, time points where the adjusted 95% bootstrap confidence interval for the difference between cue signaling the valued outcome and the cue signaling the devalued outcome excludes zero (10-sample cluster minimum); Perm, time points for which cue signaling the valued outcome is significantly (*P* < 0.05) different from cue signaling the devalued outcome from permutation test. **(t)** Area under the BLA GRABDA Z-scored curve (AUC). 2-way repeated measures ANOVA, CS x Value: F2, 18 = 6.26, *P* = 0.009; CS: F1, 18 = 10.06, *P* = 0.001; Value: F1, 9 = 5.94, *P* = 0.04. **(u)** Mediation analysis. Outcome value (devalued→valued) effect on cue-evoked dopamine: β = 73.13, SE = 17.71, t78 = 4.13, P = 0.001, 95% CI 37.87 – 108.38; effect of cue-evoked dopamine on food-port entries: β = 0.013, SE = 0.005, t78 = 2.35, P = 0.02, 95% CI 0.002 – 0.02; Direct effect of Value on entries: β = 0.75, SE = 0.91, t78 = 0.82, P = 0.41, 95% CI −1.06 – 2.57; Indirect effect of Value on entries mediated by dopamine: β = 0.92, BootSE = 0.51, BootCI 0.18 – 2.12. **(v-x)** Linear regression model using convolved Valued and Devalued CS onset, offset, and food-port entries to predict GRABDA signal at each time point 10 s prior to and 40 s after each cue onset from the event kernels. Average regression coefficient of each kernel around cue presentation (v), β coefficients from model (w; 1-way repeated measures ANOVA, CS/Value: F4,36 = 3.58, *P* = 0.01), and percentage of variance explained by each event kernel (x; 1-way repeated measures ANOVA, CS/Value: F4,36 = 2.65, *P* = 0.049). Devaluation, GRABDA: *N* = 10 (6 male); GFP: *N* = 4 (2 male). Data presented as trial-averaged, between-subject mean ± s.e.m. with individual data points. *P<0.05, **P<0.01, ***P<0.001 Bonferroni-corrected post-hoc comparisons.

We used fiber photometry to record fluorescent activity of the G-protein-coupled receptor-activation-based dopamine sensor (GRAB_DA_) in the BLA during the PIT test (Figure 1a-b, f). In separate subjects, we recorded GFP florescence. Cue onset triggered robust, transient dopamine release (Figure 1j-k, see Extended Data Figure 1-1 for non-subtracted GRAB_DA_ fluorescence response). We also detected a transient dopamine response to cue offset (Figure 1j-k). VTA_DA_ axons in the BLA were similarly activated by reward cues (Extended Data Figure 1-2). Because lever pressing and food-port entries can freely occur during cue presentation, we used a linear regression model^49, 50^ to parse the extent to which dopamine fluctuations were explained by the cues vs. concomitant behaviors. The model included kernels for cue onset and offset, food-port entries, presses on the lever earning the same outcome as the current cue, and presses on the lever earning the different outcome. Dopamine release was most heavily influenced by cue onset, indicating that the change in the reward prediction state signaled by cue onset, rather than behaviors occurring at this time, regulates dopamine release (Figure 1l-n). Thus, reward-predictive cues evoke phasic dopamine release in the BLA during decision making.

#### Cue-evoked BLA dopamine encodes predicted reward value

We next asked whether cue-evoked BLA dopamine release conveys information about the predicted reward during adaptive decision making. To do this, rats were retrained on the Pavlovian associations and then we devalued one of the food rewards by pairing it with lithium chloride (LiCl; 6 pairings). To prevent any opportunity to cache the new value to the predictive cue, devaluation occurred in the absence of the associated cue. The devaluation was effective. Rats fully and selectively rejected the LiCl-paired food (Figure 1r). Following devaluation, rats received a probe test during which each cue was presented 4 times (without accompanying reward), with intervening cue-free baseline periods. This test assesses the subjects’ ability to represent the predicted reward and use its current value to guide cue-response decisions. If subjects are using the cues to predict and consider the specific available reward, they will attenuate food-port approach during the cue paired with the devalued reward^4, 51, 52^. Indeed, indicating adaptive behavior, rats increased entries into the food-port during the cue signaling the valued reward, but not that signaling the devalued reward (Figure 1p-q).

Cue-evoked BLA dopamine release was shaped by the value of the predicted reward. The cue-evoked BLA dopamine response was larger for the cue signaling the valued reward than that signaling the devalued reward (Figure 1s-t). This adaptation of the cue-evoked dopamine response occurred without opportunity to experience the cue-reward association following devaluation and, thereby, cache the new value to the cue. Thus, via an internal model of cue-reward associations, cue-evoked BLA dopamine release can encode the value of the predicted reward. We again used a linear regression model to parse the extent to which dopamine fluctuations were explained by the cues vs. concomitant behaviors. The model included kernels for onset and offset of the cue signaling the valued reward, onset and offset of the cue signaling the devalued reward, and food-port entries. Dopamine release was most heavily influenced by cue onset, indicating that the differential dopamine response to the cues was due to differences in predicted reward value rather than behavioral responses (Figure 1v-x). The influence of cue onset on dopamine was stronger for the cue predicting the valued reward than the cue signaling the devalued reward, further indicating that cue-evoked BLA dopamine release is shaped by the value of the predicted reward. We next used mediation analysis to ask whether cue-evoked BLA dopamine mediates the influence the cue-reward prediction on adaptive behavior (Figure 1u). Predicted reward value had a significant positive influence on the magnitude of cue-evoked BLA dopamine and cue-evoked dopamine significantly positively predicted food-port entries. When dopamine was accounted for, predicted value no longer had a direct effect on entries, indicating that the magnitude of cue-evoked dopamine explains a meaningful portion (∼55%) of the influence of predicted reward value on adaptive behavior. Thus, predictive cues trigger dopamine release in the BLA, this reflects the value of the predicted reward, and is associated with the ability to use cue-reward predictions for adaptive decision making.

### Cue-evoked VTA_DA_→BLA activity mediates the influence of reward predictions on decision making

#### Cue-evoked VTA_DA_→BLA projection activity mediates action selection

We next asked whether cue-evoked VTA_DA_→BLA activity mediates the influence of cue-reward predictions on decision making. To allow for transient inactivation of VTA_DA_ axons and terminals in the BLA, we cre-dependently expressed the inhibitory opsin archaerhodopsin T (ArchT) or tdTomato control bilaterally in VTA_DA_ neurons of male and female tyrosine hydroxylase (Th)-cre rats and implanted optical fibers bilaterally over BLA (Figure 2a-b). Rats first received instrumental conditioning, without manipulation, in which each of two different lever-press actions earned one of two distinct food rewards (Figure 2c-d). Rats then received Pavlovian conditioning, also without manipulation, during which 2 distinct 30-s auditory cues each predicted delivery of one of the unique food rewards at cue offset (Figure 2c, e). They were then given a PIT test during which the levers were available and each cue was presented 4 times in pseudorandom order to assess its influence over action selection. We optically inhibited (532 nm, 10 mW, 2.5 s) VTA_DA_ axons and terminals in the BLA coincident with each cue onset during this test. We restricted optical inhibition to cue onset to inhibit only the phasic cue onset-induced dopamine elevation. We reasoned that, if cue-evoked VTA_DA_→BLA activity supports the reward predictions that enable action selection, inhibiting this signal should prevent cues from biasing action selection. Indeed, inhibition of cue-evoked VTA_DA_→BLA activity attenuated the ability of the cues to bias action selection towards the lever associated with the same predicted reward than that associated with the different reward (Figure 2g-h). Inhibition of cue-evoked VTA_DA_→BLA activity did not disrupt expression of the conditional approach responses to the shared goal location (Figure 2i), a behavior that does not require prediction of the identity of the specific reward. Providing further evidence that BLA dopamine supports cue-reward predictions, we found that pharmacological inactivation of BLA dopamine D1 receptors similarly disrupted the ability of cues to bias action selection without affecting conditional approach responses (Extended Data Figure 2-2a-i). Thus, cue-evoked VTA_DA_→BLA activity mediates the influence of the cues on action selection, but not general motivated behavior.

**Figure 2.**
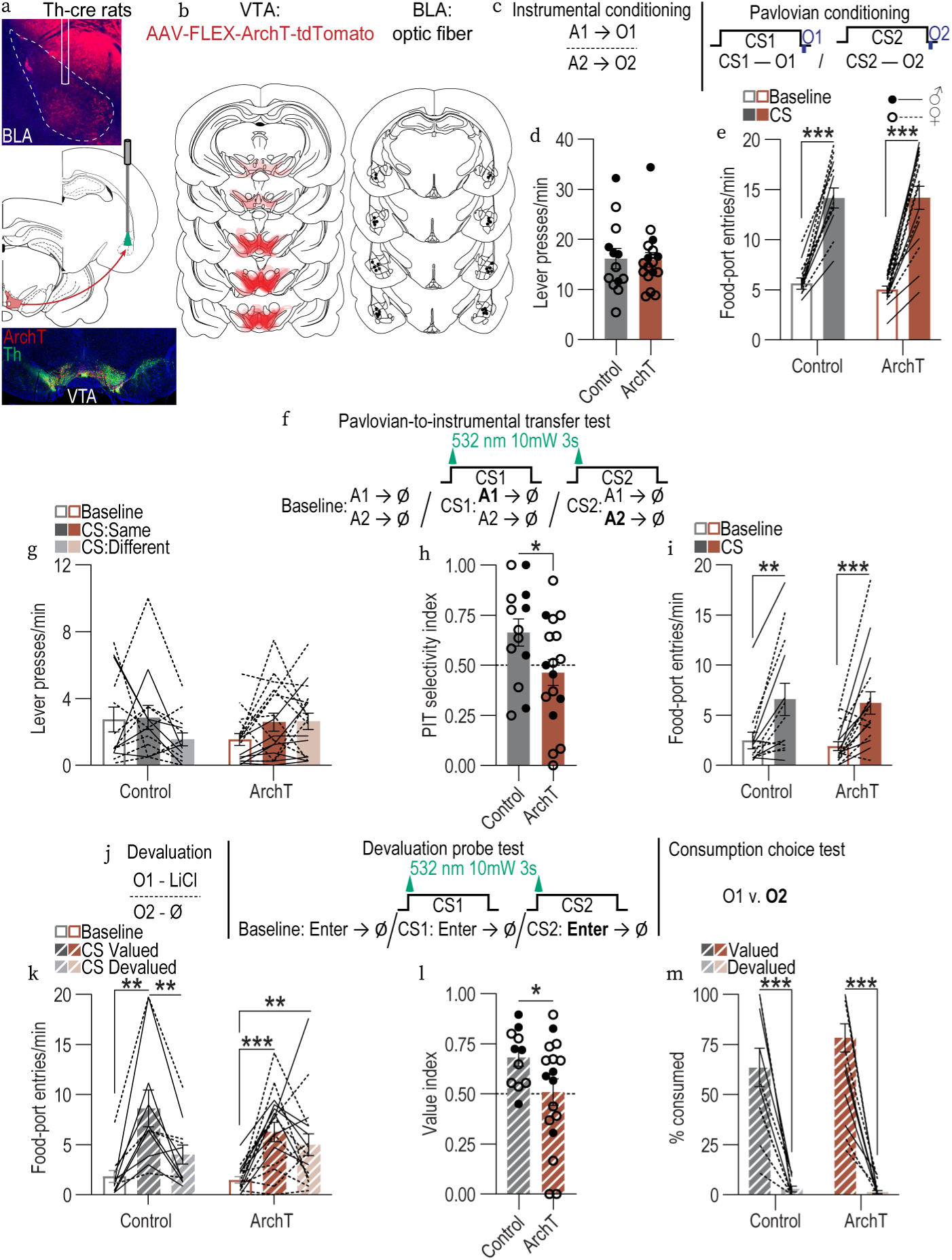
Cue-evoked VTA_DA_→BLA activity mediates the influence of cue-reward predictions on decision making. **(a)** Bottom: Representative fluorescent image of ArchT-tdTomato expression in VTADA neurons. Middle: Strategy for bilateral optogenetic inhibition of VTA_DA_→BLA projections. Top: Representative image of fiber placement in the vicinity of immunofluorescent ArchT-tdTomato-expressing VTA_DA_ axons and terminals in BLA. **(b)** Schematic representation of ArchT-tdTomato expression in VTA and placement of optical fiber tips in BLA for all subjects. **(c-e)** Instrumental and Pavlovian conditioning **(c)** Conditioning procedures. A, action (left or right lever press); CS, 30-s conditioned stimulus (aka, “cue”, white noise or click) followed immediately by reward outcome (O, ensure solution or grain pellet). **(d)** Lever-press rate averaged across levers and across the final 2 instrumental sessions. 2-tailed students t-test with Welch’s correction, t23.02 = 0.10, *P* = 0.92, 95% confidence interval (CI) −5.24 - 4.93. **(e)** Food-port entry rate during averaged across cues and across the final 2 Pavlovian conditioning sessions. 2-way repeated measures ANOVA, CS: F1, 28 = 2.04, *P* = 0.03; Virus: F1, 28 = 0.09, *P* = 0.77; CS x Virus: F1, 28 = 0.19, *P* = 0.66. **(f-i)** Outcome-specific Pavlovian-to-instrumental transfer test. **(f)** PIT procedure. Ø, no reward was delivered. **(g)** Lever-press rates on the lever that earned the “Same” outcome as predicted or current cue or on the other available lever (Different) compared to pre-cue baseline press rate averaged across levers. 2-way repeated measures ANOVA, CS/Lever x Virus: F2,56 = 3.28, *P* = 0.045; CS/Lever: F2,56 = 1.22, *P* = 0.30; Virus: F1, 28 = 0.05, *P* = 0.82. **(h)** PIT selectivity index [Same presses/(Same + Different presses)]. 2-tailed students t-test with Welch’s correction, t26.90 = 2.13, *P* = 0.04, 95% CI −0.39 - −0.007. **(i)** Food-port entry rate. 2-way repeated measures ANOVA, CS: F1, 28 = 30.73, *P* < 0.0001; Virus: F1, 28 = 0.13, *P* = 0.72; CS x Virus: F1, 28 = 0.02, *P* = 0.90. PIT: Control *N* = 13 (7 male); ArchT *N* = 17 (7 male). **(j-m)** Outcome-specific devaluation test. **(j)** Devaluation procedures. LiCl, lithium chloride 0.3M, 1.5% volume/weight. **(k)** Food-port rates during the 35-s pre-cue baseline periods compared to that during the cue (30-s cue periods + 5-s post-cue periods) predicting the non-devalued (Valued) and Devalued outcome. 2-way repeated measures ANOVA, CS: F1.80, 46.76 = 25.55, *P* < 0.0001; Virus: F1, 26 = 0.26, *P* = 0.62; CS x Virus: F1.80, 46.76 = 2.12, *P* = 0.14. **(l)** Value index [Valued CS entries/(Valued + Devalued CS entries)]. 2-tailed students t-test with Welch’s correction, t25.24 = 2.20, *P* = 0.04, 95% confidence interval (CI) −0.33 - −0.01. **(m)** Post-devaluation consumption test. % of total available food (10 g or 10 mL) consumed during 10-min test. 2-way repeated measures ANOVA, Value: F1, 26 = 134, *P* < 0.0001; Virus: F1, 26 = 1.32, *P* = 0.26; Value x Virus: F1, 26 = 1.89, *P* = 0.18. Devaluation: Control *N* = 11 (7 male); ArchT *N* = 17 (7 male). Data presented as trial-averaged, between-subject mean ± s.e.m. with individual data points. *P<0.05, **P<0.01, ***P<0.001 Bonferroni-corrected post-hoc comparisons.

#### Cue-evoked VTA_DA_→BLA projection activity mediates sensitivity to outcome devaluation

To provide converging evidence that cue-evoked VTA_DA_→BLA activity supports the influence of cue-reward predictions on adaptive decision making, we next asked whether cue-evoked VTA_DA_→BLA activity mediates the ability to adapt cue responses following devaluation of the predicted reward (Figure 2j). Rats were retrained on the Pavlovian associations and then we devalued one of the food rewards by pairing it with LiCl (6 pairings) in the absence of the associated cue. The devaluation was effective (Figure 2m). Following devaluation, rats received a probe test during which each cue was presented (without reward). We optically inhibited (532 nm, 10 mW, 2.5 s) VTA_DA_ axons and terminals in the BLA coincident with each cue onset during this test. Because cue-evoked BLA dopamine encodes the value of the predicted reward, we reasoned that inhibiting this signal would prevent subjects from adapting their cue responses based on the predicted reward’s current value. Indeed, whereas controls increased entries into the food-port during the cue signaling the valued reward more than that signaling the devalued reward, inhibition of cue-evoked VTA_DA_→BLA activity caused elevated approach responses during both cues, thereby attenuating sensitivity of the conditional response to devaluation (Figure 2k-l; see also Extended Data Figure 2-1). Thus, cue-evoked VTA_DA_→BLA activity mediates the ability to adapt cue-response decisions based on the value of the predicted reward. Together, these data indicate that cue-evoked BLA dopamine supports the influence of identity-specific cue-reward predictions on adaptive decision making.

### Cue-evoked VTA_DA_→BLA activity mediates the constraining influence of reward predictions on learning

#### Cue-evoked VTA_DA_→BLA projection activity mediates blocking

If cue-evoked VTA_DA_→BLA activity supports reward prediction, it should not only affect decision making, but should also influence new learning. Learning is regulated by predictions. We use information available from environmental cues to generate predictions about forthcoming events and tend to learn when our expectations are violated^7–9^. When our predictions are fully realized, learning about new, redundant predictors is blocked. Therefore, we reasoned that if cue-evoked VTA_DA_→BLA activity supports reward predictions, attenuating this signal should prevent such predictions and unblock learning. To test this and, thereby, provide more evidence for a function in reward prediction, we inhibited cue-evoked VTA_DA_→BLA activity during a classic Kamin blocking procedure^7^ in which visual cues that already reliably predict particular rewards block formation of associations between novel auditory cues and those specific rewards^16, 53^. We first cre-dependently expressed ArchT bilaterally in VTA_DA_ neurons of male and female Th-cre rats and implanted optical fibers bilaterally over BLA (Figure 3a-b). For controls, we either cre-dependently expressed tdTomato bilaterally in VTA_DA_ neurons of Th-cre rats or infused an ArchT AAV in Th-cre- rats and, in both cases, implanted optical fibers bilaterally over BLA. Rats first received instrumental conditioning, without manipulation, in which each of two different lever-press actions earned one of two distinct food rewards (Figure 3c-d). They then received visual cue Pavlovian conditioning, also manipulation-free, during which each of two, distinct, 30-s visual cues were paired with a unique food outcome at cue offset (e.g., house light—ensure/flashing light—pellet). Both groups developed Pavlovian conditional goal-approach responses to the visual cues (Figure 3e). Rats next received compound conditioning (aka blocking) during which each of the visual cues was presented concurrent with an auditory cue for 30 s terminating in the delivery of the same outcome already associated with the visual cue (e.g., house light + white noise—ensure/flashing light + click—pellet; Figure 3f). During each compound conditioning session, we optically inhibited (532 nm, 10 mW, 2.5 s) VTA_DA_ axons and terminals in the BLA coincident with each cue onset. Inhibition of cue-evoked VTA_DA_→BLA activity did not affect conditional goal-approach responses to the cues (Figure 3g). During the compound cues, both groups increased entries into the food-delivery port. To assess encoding of identity-specific cue-reward associations, we next gave rats a PIT test with only the auditory cues, without manipulation. Controls showed evidence of blocking: the auditory cues were not capable of guiding choice behavior. Inhibition of cue-evoked VTA_DA_→BLA activity during compound training did, however, enable the encoding of identity-specific cue-reward associations. Rats in this group were able to use the auditory cues to know which specific outcome was predicted to selectively increase presses on the lever associated with that same reward (Figure 3i-j). Thus, inhibition of cue-evoked VTA_DA_→BLA activity unblocked learning of the identity-specific cue-reward associations that enable subsequent reward predictions. Inhibition of cue-evoked VTA_DA_→BLA activity during compound training also caused subjects to subsequently show more conditional goal-approach responses. Therefore, cue-evoked VTA_DA_→BLA activity mediates the ability of cue-reward predictions to constrain new learning.

**Figure 3.**
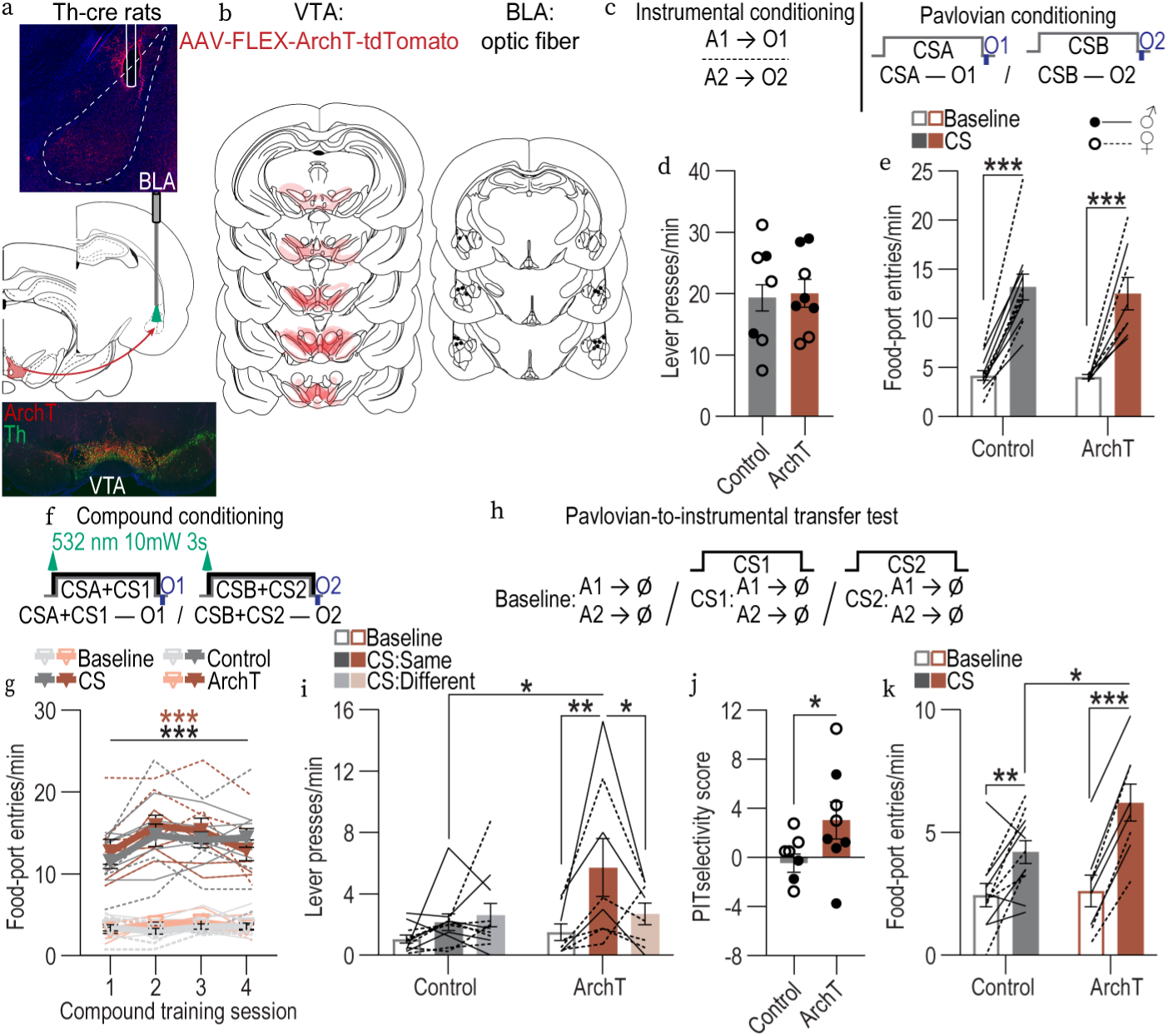
Cue-evoked VTA_DA_→BLA activity mediates the ability of cue-reward predictions to constrain learning. **(a)** Bottom: Representative fluorescent image of ArchT-tdTomato expression in VTA_DA_ neurons. Middle: Strategy for bilateral optogenetic inhibition of VTA_DA_→BLA projections. Top: Representative image of fiber placement in the vicinity of immunofluorescent ArchT-tdTomato-expressing VTA_DA_ axons and terminals in BLA. **(b)** Schematic representation of ArchT-tdTomato expression in VTA and placement of optical fiber tips in BLA for all subjects. **(c-e)** Instrumental and visual cue Pavlovian conditioning **(c)** Conditioning procedures. A, action (left or right lever press); CSA/B, 30-s conditioned stimulus (aka, “cue”, white house light or 2hz flashing stimulus lights) followed immediately by reward outcome (O, ensure solution or grain pellet). **(d)** Lever-press rate averaged across levers and across the final 2 instrumental sessions. 2-tailed students t-test with Welch’s correction, t16.09 = 0.23, *P* = 0.82, 95% confidence interval (CI) −5.93 – 7.34. **(e)** Food-port entry rate during averaged across cues and across the final 2 Pavlovian conditioning sessions. 2-way repeated measures ANOVA, CS: F1,17 = 99.94, *P* < 0.0001; Virus: F1,17 = 0.10, *P* = 0.75; CS x Virus: F1,17 = 0.10, *P* = 0.76. **(f-g)** Compound conditioning (aka, “blocking”). **(f)** Compound conditioning procedure. CSA/B: house light or flashing lights; CS1/CS2: white noise or click followed immediately by reward outcome (O, ensure solution or grain pellet). **(g)** Food-port entry rate. 3-way repeated measures ANOVA, Training x CS: F3,51 = 10.10, *P* < 0.0001; Compound training: F3,51 = 6.91, *P* = 0.0005; CS: F1,17 = 179.3, *P* < 0.0001; Virus: F1,17 = 0.19, *P* = 0.67; Training x Virus: F3,51 = 2.03, *P* = 0.12; Virus x CS: F1,17 = 0.02, *P* = 0.89; Training x Virus x CS: F3,51 = 2.13, *P* = 0.11. **(h-k)** Auditory Pavlovian-to-instrumental transfer test. **(h)** Auditory cue PIT procedure. Ø, no reward was delivered. **(g)** Lever-press rates on the lever that earned the “Same” outcome as predicted or current cue or on the other available lever (Different) compared to pre-cue baseline press rate averaged across levers. 2-way repeated measures ANOVA, CS/Lever: F2,34 = 3.92, *P* = 0.03; Virus: F1,17 = 2.22, *P* = 0.15; CS/Lever x Virus: F2,34 = 7.55, *P* = 0.002. **(h)** PIT selectivity index [Same presses/(Same + Different presses)]. 2-tailed students t-test, t17 = 2.25, *P* = 0.04, 95% CI 0.21 – 6.76. **(i)** Food-port entry rate. 2-way repeated measures ANOVA, CS x Virus: F1,17 = 8.18, *P* = 0.01; CS: F1,17 = 68.67, *P* < 0.0001; Virus: F1,17 = 2.13, *P* = 0.16. Control *N* = 11 (6 male, 3 Cre- 8 Cre +); ArchT *N* = 10 (5 male). Data presented as trial-averaged, between-subject mean ± s.e.m. with individual data points. *P<0.05, **P<0.01, ***P<0.001 Bonferroni-corrected post-hoc comparisons.

## DISCUSSION

Here we explored the function of BLA dopamine in reward prediction. We found that reward-predictive cues trigger dopamine release in the BLA during decision making and, correspondingly, cue-evoked VTA_DA_→BLA activity and BLA dopamine D1 receptors mediate the influence of cue-reward predictions to bias action selection. Cue-evoked BLA dopamine release encodes the value of the predicted reward and cue-evoked VTA_DA_→BLA activity mediates the ability to adapt cue-response decisions based on such value. Lastly, we found that cue-evoked VTA_DA_→BLA activity mediates the constraining influence of cue-reward predictions on learning. Thus, reward-predictive cues evoke dopamine release in the BLA to support the reward predictions that both enable adaptive decision making and constrain learning.

Cue-evoked BLA dopamine mediates the influence of cue-reward prediction on decision making. We found that reward-predictive cues trigger dopamine release in the BLA, consistent with the activity of midbrain dopamine neurons^23–25^ and prior evidence that reward cues come to evoke BLA dopamine release with learning^16, 46^. We add that this cue-evoked dopamine release occurs transiently at cue transition points (onset and offset) during decision making. Thus, cue-evoked dopamine transients are positioned to support a transition in the reward-prediction state. Correspondingly, inhibiting either transient cue-evoked VTA_DA_→BLA activity or BLA dopamine D1 receptors disrupted the ability to use the cue-reward predictions to guide action selection. It did not, however, affect performance of general motivated behavior that does not require a detailed reward prediction. Thus, cues evoke dopamine release in the BLA to support the ability to use the specific predictions those cues generate to guide decision making.

Cue-evoked VTA_DA_→BLA activity mediates the influence of predicted reward value on decision making. We found that BLA dopamine release encodes the model-based value of the predicted reward. Cue-evoked dopamine was lower when the predicted reward had been devalued. This encoding of predicted reward value occurred despite no prior opportunity to experience the cue-reward association following devaluation and, thereby, cache the new value to the cue. Thus, rather than encoding the cached value of the reward learned during training, cue-evoked BLA dopamine is shaped by a dynamic valuation process to reflect the current motivational value of the predicted reward. This could have occurred during via mediated conditioning whereby, following pairing with illness, reward experience triggered a mental representation of the predictive cue allowing that cue to become linked to the new reward value and, thereby, subsequently trigger less dopamine release. Other possibility is that the cue-evoked dopamine could have been shaped online during the test by the cue-triggered reward prediction. In either case, information derived from an internal model of environmental cue-reward associations shapes cue-evoked BLA dopamine. Importantly, this encoding of predicted reward value was behaviorally meaningful, explaining a significant portion of the ability of the cues to enable adaptive behavior. Correspondingly, inhibiting transient cue-evoked VTA_DA_→BLA activity disrupted the ability to use the cue-reward predictions to adapt behavior based on the current value of the predicted reward. Prediction of a devalued outcome caused the cues to elicit less dopamine and this was associated with lower cue responses, yet, interestingly, inhibiting cue-evoked VTA_DA_→BLA activity caused higher responses to the cue signaling the devalued reward. Thus, inhibition of cue-evoked VTA_DA_→BLA activity may have caused the conditional goal approach responses to become controlled by system that relies on the previously cached value of the cue, rather than the current value inferred by the internal cue-reward model. Indeed, there are multiple learning and decision systems that rely on different forms of value^54, 55^. Despite only brief inhibition at cue onset, the effects on decision making persisted during and just after the cue further supporting the notion that cue-evoked dopamine transients support a transition in the reward-prediction state^56^ that controls subsequent adaptive decision making.

Cue-evoked VTA_DA_→BLA activity mediates the suppressive influence of cue-reward predictions on learning. In addition to influencing decision making, strong cue-reward predictions can also constrain learning. We found that preventing cues from activating VTA_DA_→BLA projections removed the suppressive influence of reward predictions on new learning, unblocking the encoding of identity-specific cue-reward associations. Thus, cue-evoked VTA_DA_→BLA activity supports reward predictions to constrain new learning. This finding contrasts with prior evidence that optogenetically shunting reward cue-evoked VTA_DA_ neuron activity does *not* unblock learning^15^. This discrepancy is likely explained by one or both of two factors. First, the prior work used only Pavlovian conditional food-port approach responses to assess learning and did not include tests of the content of learning. We used a PIT test to ask whether identity-specific cue-reward learning was unblocked to subsequently guide instrumental decision making. Second, whereas the prior work non-selectively inhibited VTA_DA_ neurons, we specifically inhibited VTA_DA_→BLA activity. Thus, VTA_DA_ neuron function may differ based on pathway, with the VTA_DA_→BLA projection supporting reward predictions.

Findings from this study have important implications for how we conceptualize dopamine function. One prominent theory proposes that dopamine is a teaching signal to support predictive learning^10–13^. Often viewed as a competing account, dopamine is also theorized to function online to support motivated behavior^19–22^. But these theories need not be incompatible^19^. Via projections to the BLA, dopamine neurons were recently discovered to support cue-reward learning^16^. Rewards trigger dopamine release in the BLA and this functions as a teaching signal to help link the reward identity to a predictive cue so that cue can later generate the predictions needed for decision making^16^. With learning, reward-predictive cues come to evoked BLA dopamine release^16^. Here we discovered that this cue-evoked BLA dopamine encodes information about the predicted reward and supports reward prediction. Thus, dopamine can support both learning and prediction. Reward-evoked BLA dopamine supports predictive learning and cue-evoked BLA dopamine supports prediction. Whether cue-evoked BLA dopamine also mediates higher-order learning, additional to its function in reward prediction, is a ripe question for future investigation.

These findings open the door to several important future questions. One concerns the mechanisms through which cue-evoked VTA_DA_→BLA neuron activity mediates reward prediction. Our pharmacology data indicate this function is mediated, at least in part, via activation of dopamine D1 receptors. D1 receptors are expressed on both BLA principal neurons and interneurons^57^ where they can positively modulate projection neuron excitability and also enhance interneuron-driven inhibition, thereby modulating network gain^58–61^. Thus, VTA_DA_ input is well positioned to regulate how reward-predictive cues influence BLA circuit activity. Such activity is likely to include BLA projections to the lateral^62^ and medial^63^ orbitofrontal cortex, which mediate influence of cue-reward predictions on decision making. Our pharmacological results implicate dopamine release in supporting reward prediction, but an important question is whether glutamate corelease^61, 64^ also contributes. Another new question is how VTA_DA_→BLA function in cue-reward prediction differs from the function of other VTA_DA_ pathways. For example, NAc core dopamine is also shaped by cue-reward predictions and regulates their influence on decision making^65, 66^, but this may reflect distinct components of predictive processing. Finally, it will be important to determine the mechanisms that allow BLA dopamine to support both learning and decision making.

Here we show that the BLA dopamine is evoked by reward-predictive cues to support the predictions critical for adaptive learning and decision making. Disruptions to reward prediction can lead to ill-informed decisions and unregulated learning. This is characteristic of the cognitive symptoms underlying many psychiatric diseases. Thus, these data may also aid our understanding and treatment of substance use disorder and mental health conditions marked by disruptions to dopamine, learning, and decision making.

## METHODS

### Subjects

Male and female wildtype Long-Evans rats and transgenic Long-Evans rats expressing Cre recombinase under control of the tyrosine hydroxylase promoter (Th-cre) aged 8 - 12 weeks at the time of surgery served as subjects. Wild type rats were either sourced from our internal colony or purchased from Inotiv (HsdBlu:LE, West Lafayette, ID). Rats were housed in a temperature (68-79°F) and humidity (30-70%) regulated vivarium. They were initially housed in same-sex pairs and then following surgery housed individually to preserve implants. Rats were provided with water *ad libitum* in the home cage and were maintained on a food-restricted 12-18 g daily diet (Lab Diet, St. Louis, MO) to maintain approximately 85-90% free-feeding body weight. Rats were handled for 3-5 days prior to the onset of each experiment. Separate groups of naïve rats were used for each experiment. Experiments were performed during the dark phase of a 12:12 hr reverse dark/light cycle (lights off at 7AM). All procedures were conducted in accordance with the NIH Guide for the Care and Use of Laboratory Animals and were approved by the UCLA Institutional Animal Care and Use Committee.

### Surgery

We used standard surgical procedures described previously^16, 36, 63, 67, 68^. Rats were anesthetized with isoflurane (4–5% induction, 1–2% maintenance), and a nonsteroidal anti-inflammatory agent was administered pre- and postoperatively to minimize pain and discomfort. Surgical details for each experiment are described below. In all cases, surgery occurred prior to the onset of behavioral training.

### Behavioral procedures

#### Apparatus

Training took place in Med Associates conditioning chambers (East Fairfield, VT) housed within sound- and light-attenuating boxes, described previously^38^. Each chamber had grid floors and contained 2 retractable levers that could be inserted to the left and right of a recessed food-delivery port (magazine) on the front wall. A photobeam entry detector was positioned at the entry to the food port. Each chamber was equipped with a syringe pump to deliver 60% vanilla-flavored ensure^®^ solution (Abbott, Abbott Park, IL) in 0.1 ml increments through a stainless-steel tube into one well of the food port and a pellet dispenser to deliver 45-mg food pellets (Bio-Serv, Frenchtown, NJ) into another well of the same port. A white noise generator was attached to a speaker on the wall opposite the levers and food-delivery port. A clicker was also mounted on this wall. Stimulus lights were positioned above each lever. A 3-watt, 24-volt house light mounted on the top of the back wall opposite the food port provided illumination, except in Pavlovian blocking experiments for which it was used as a conditioned stimulus. A fan mounted to the outer chamber provided ventilation and external noise reduction. For fiber photometry experiments, chambers were outfitted a Pigtailed 1×1 Fiber-optic Rotary Joint Gen.2 (Doric Lenses, Quebec, Canada) connecting the fiber optic patch cord from the fiber photometry system (MBF Bioscience, Williston, VT) to the rat to allow free movement in the chamber. For optogenetic manipulations, chambers were outfitted with an Intensity Division Fiberoptic Rotary Joint (Doric Lenses, Quebec, QC, Canada) connecting the output fiber optic patch cords to a laser (CNI Laser, ChangChun, JiLin, China) positioned outside of the chamber.

##### Pavlovian delay conditioning with Pavlovian-to-instrumental transfer and outcome-specific devaluation tests

###### Magazine conditioning

Rats first received 2 days of training to learn where to receive the ensure (60%, vanilla, 0.1 ml/delivery) and food pellet (45 mg grain) outcomes. Each day included 2 sessions, separated by approximately 1 hr, order counterbalanced across days, one with 30 non-contingent deliveries of ensure and one with 30 grain pellet deliveries (60-s intertrial interval, ITI).

###### Instrumental conditioning

Rats next received 11 days, minimum, of instrumental conditioning. Each day consisted of 2 training sessions, one with the left lever and one with the right lever, separated by at least 1 hr with order alternated across days. Each action was reinforced with one of the different food outcomes (e.g., left press→grain pellets/right press→ensure solution). Lever-outcome pairings were counterbalanced at the start of the experiment within each group. Each session terminated after 20 outcomes had been earned or 30 min had elapsed. Actions were continuously reinforced on the first day and then escalated ultimately to a random-ratio (RR) 10 schedule of reinforcement in which a variable number of presses (average = 10) were required to earn a reward.

###### Pavlovian conditioning

All rats received 6 sessions of Pavlovian conditioning (1 session/day on consecutive days) to learn to associate each of 2 auditory cues (aka conditioned stimuli; 80-82 db), click (10 Hz) and white noise, with a specific food outcome, ensure solution or grain pellets. Each 30-s cue terminated with the delivery of its associated outcome. For half the subjects, click terminated in the delivery of ensure and noise predicted pellets, with the other half receiving the opposite arrangement. Each session consisted of 8 click and 8 white noise presentations. Cues were delivered pseudo-randomly with a variable 1.5 – 3-min ITI (mean = 2.5 min).

###### Instrumental retraining and extinction

Following Pavlovian conditioning, rats received one instrumental retraining session on the RR-10 reinforcement schedule. Rats then received one session of instrumental extinction to establish a low level of pressing. During this single 20-min session both levers were available but pressing was not reinforced.

###### Outcome-specific Pavlovian-to-instrumental transfer tests

Rats next received an outcome-specific Pavlovian-to-instrumental transfer (PIT) test. During the PIT test, both levers were continuously present, but pressing was not reinforced. After 5 min of lever-pressing extinction, each 30-s cue was presented separately 4 times, separated by a fixed 2.5-min ITI, in alternating order. Cue order was counterbalanced across subjects. No outcomes were delivered following cue presentation.

###### Pavlovian retraining

Following the PIT test, to reestablish the Pavlovian associations, rats received 2 Pavlovian conditioning sessions, identical to those above.

###### Outcome-specific devaluation by conditioned taste aversion

Following retraining, one of the food rewards was devalued by pairing with the malaise-inducing agent lithium chloride (LiCl). In the conditioning chambers, rats were given 30, non-contingent deliveries of one reward type (60-s intertrial interval, ITI) followed immediately by a i.p. injection of LiCl (0.3M, 1.5% volume/weight). For the control, rats were given 30, non-contingent deliveries of the other reward type (60-s intertrial interval, ITI) in the conditioning chamber, without subsequent LiCl injection. Rats received 1 session/day with 12 total sessions (6 devaluation and 6 control) in the order 3 devaluation, 3 control, 2 devaluation, 3 control and 1 devaluation.

###### Outcome-specific devaluation probe test

24 hr after the last session, rats next received an outcome-specific devaluation probe test. Each 30-s cue was presented separately 8 times, separated by a variable 2.5-min ITI, in alternating order. Cue order was counterbalanced across subjects. No outcomes were delivered following cue presentation.

###### Outcome-specific devaluation consumption test

After the devaluation probe test, rats received a consumption choice test to confirm the efficacy of the conditioned taste aversion. Rats were given access to 10 g of pellets and 10 mL of ensure solution and allowed to consume freely for 10 min.

##### Outcome-specific blocking and Pavlovian-to-instrumental transfer

###### Magazine conditioning

Rats first received 2 days of training to learn where to receive the ensure (60%, vanilla 0.1 ml/delivery) and food pellet (45 mg grain) rewards. Each day included 2 sessions, separated by approximately 1 hr, order counterbalanced across days, one with 30 non-contingent deliveries of ensure and one with 30 grain pellet deliveries (60-s ITI). The house light was off during these sessions.

###### Instrumental conditioning

Rats next received 11 days, minimum, of instrumental conditioning. Each day consisted of 2 training sessions, one with the left lever and one with the right lever, separated by at least 1 hr with order alternated across days. Each action was reinforced with one of the different food outcomes (e.g., left press→grain pellets/right press→ensure solution). Lever-outcome pairings were counterbalanced at the start of the experiment within each group. Each session terminated after 20 outcomes had been earned or 30 min had elapsed. Actions were continuously reinforced on the first day and then escalated ultimately to a RR-10 schedule of reinforcement.

###### Pavlovian conditioning

Rats received 12 sessions of visual cue Pavlovian conditioning (1 session/day on consecutive days) in a dark operant chamber to learn to associate visual cues with the food outcomes. Each of 2 30-s visual cues, house light or flashing stimulus lights (2 hz), was paired with a specific food outcome, ensure (60%, vanilla, 0.1 ml/delivery) or grain pellets (e.g., house light—ensure solution/flashing light—pellet). Cue-reward pairings were counterbalanced within groups and with respect to instrumental lever-outcome pairings. For half the subjects, the house light terminated in the delivery of ensure and flashing lights predicted pellets, with the other half receiving the opposite arrangement. Each session consisted of 16 house light and 16 flashing light presentations. Cues were delivered pseudo-randomly with a variable 1.5 – 3-min ITI (mean = 2.5 min).

###### Instrumental retraining and extinction

Following Pavlovian conditioning, rats received one instrumental retraining session on the RR-10 reinforcement schedule. Rats then received one session of instrumental extinction to establish a low level of pressing. During this single 20-min session both levers were available but pressing was not reinforced.

###### Preexposure

Rats received one day of preexposure to the auditory stimuli. Click and noise were independently presented pseudo-randomly for 30-s, 8 times each with a variable 1.5 – 3-min ITI (mean = 2.5 min).

###### Compound conditioning

Rats next received 4 compound conditioning sessions (1 session/day on consecutive days) in which the house light and flashing stimulus light cues were each presented in compound with a distinct auditory stimulus, click (10 Hz) or white noise (80-82 dB). For half the subjects in each group, the house light was presented simultaneously for 30 s with the click and the flashing lights concurrent noise for 30 s. The other half of subjects received the opposite arrangement. Visual-auditory cue pairings were counterbalanced within groups and with respect to instrumental and visual cue-reward contingencies. Each compound stimulus terminated in the reward paired with the visual stimulus during initial Pavlovian conditioning (e.g., house light + white noise—ensure solution/flashing light + click—pellet). Each compound conditioning session consisted of 8, 30-s presentations of each compound cue, terminating in the delivery of the associated food outcome. Compound cues were delivered pseudo-randomly with a variable 1.5 – 3-min ITI (mean = 2.5 min).

###### Outcome-specific Pavlovian-to-instrumental transfer tests

Rats next received an outcome-specific PIT test. During the PIT test, both levers were continuously present, but pressing was not reinforced. After 5 min of lever-pressing extinction, each 30-s cue was presented separately 4 times, separated by a fixed 2.5-min ITI, in alternating order. Cue order was counterbalanced across subjects. No outcomes were delivered following cue presentation. The house light was off at test.

##### Data collection

Entries into the food-delivery port and/or lever presses were recorded continuously for each session.

### Fiber photometry recordings of dopamine release in the BLA during Pavlovian-to-instrumental transfer and outcome-specific devaluation tests

#### Subjects

Eleven male (*N* = 7) and female (*N* = 4) Long Evans rats (Th-cre- littermates, *N* = 6; Inotiv, *N* = 5) aged 8-12 weeks at the time of surgery were used to record dopamine release in the BLA during the Pavlovian-to-instrumental transfer and outcome-specific devaluation tests. Subjects without sufficient fiber photometry GRAB_DA2m_ signal of sufficient quality and/or were excluded from the dataset prior to analysis (*N* = 8). An additional 4 male (*N* = 2) and female (*N* = 2) Long Evans rats (Th-cre- littermates, *N* = 4) were used to record GFP fluorescence changes as a control. One subject was excluded from the PIT dataset due to poor quality signal (Final *N* = 10, 6 male). One subject was excluded from the devaluation dataset due to illness (Final *N* = 10, 6 male).

#### Surgery

Rats were infused unilaterally with AAV encoding the GPCR-activation-based dopamine sensor GRAB_DA2m_ (AAV9-hsyn-GRAB_DA2m, Addgene, Watertown, MA) or control fluorophore (AAV8-hSYN-GFP). Virus (0.6 µl) was infused unilaterally into the BLA (AP: −2.7; ML: ±5.0; DV: −8.6 from bregma). Optical fibers (400-µm diameter, 0.37 NA, MBF Bioscience, Williston, VT) were implanted bilaterally 0.1 mm dorsal to the infusion site. Virus was infused at a rate of 0.1 µl/min using 28-gauge injectors (Plastics One, Roanoke, VA) and injectors were left in place for 10 min. Experiments commenced approximately 2 weeks after surgery to allow sufficient expression in the BLA.

#### Fiber photometry recordings

Rats first received magazine training, instrumental and Pavlovian conditioning, as above. During the last 3 sessions of Pavlovian conditioning and instrumental retraining were attached to the optical tether, but no light was delivered. Following training, fiber photometry was used to image GRAB_DA2m_ fluorescent changes in BLA neurons during the PIT test. Rats were retrained and received conditioned taste aversion as above, and then fiber photometry was used to record image GRAB_DA2m_ fluorescent changes in BLA neurons during the outcome-specific devaluation probe test. Fiber photometry was conducted using a commercial fiber photometry system (MBF Bioscience). 470 nm excitation light was adjusted to approximately 90-100 µW at the tip of the patch cord (fiber core diameter: 400 µm; Doric Lenses). Fluorescence emission was passed through a 535 nm bandpass filter and focused onto the complementary metal-oxide semiconductor (sCMOS) ultra-sensitive camera sensor through a tube lens. Samples were collected at 20 Hz, interleaved, sequentially between 470 and 415 nm excitation channels, using a custom Bonsai^69^ workflow. Time stamps of task events were collected simultaneously through an additional synchronized camera aimed at the Med Associates interface, which sent light pulses coincident with task events. Signals were saved using Bonsai software and exported to MATLAB (MathWorks, Natick, MA) for analysis.

### Fiber photometry recordings of calcium activity in dopaminergic axons and terminals in the BLA during Pavlovian-to-instrumental transfer and outcome-specific devaluation tests

#### Subjects

Five male (*N* = 4) and female (*N* = 4) Th-Cre Long Evans rats aged 9-12 weeks at the time of surgery were used to record calcium activity in dopaminergic axons and terminals in the BLA during the Pavlovian-to-instrumental transfer test. Subjects without sufficient fiber photometry aGCamp6s signal of sufficient quality (*N* = 4) or with fibers not placed in the vicinity of GCaMP expression (N = 3) were excluded from the dataset prior to analysis.

#### Surgery

Rats were infused bilaterally with AAV encoding the axonally-targeted genetically encoded calcium indicator GCaMP6s (AAV9-FLEX-hSyn-aGcamp6s, Addgene). Virus (0.4 µl) was infused bilaterally into the VTA (AP: - 5.3; ML: ±0.75; DV: −8.35 mm from bregma). Optical fibers (400-µm diameter, 0.37 NA, MBF Bioscience) were implanted bilaterally in the BLA (AP: −2.7; ML: ±5.0; DV: −8.6 from bregma). Virus was infused at a rate of 0.1 µl/min using 28-gauge injectors and injectors were left in place for 10 min. Experiments commenced approximately 4 weeks after surgery to allow sufficient expression in the BLA.

#### Fiber photometry recordings

Rats received magazine training, instrumental and Pavlovian conditioning, as above. During these conditioning sessions rats were attached to the optical tether, but no light was delivered. Following training, fiber photometry was used to image axonal GCaMP6s fluorescent changes in BLA neurons during the Pavlovian-to-instrumental transfer test. Rats were retrained and received conditioned taste aversion as above, and then fiber photometry was used to record image axonal GCaMP6s fluorescent changes in BLA neurons during the outcome-specific devaluation probe test. Fiber photometry was conducted using a commercial fiber photometry system (MBF Bioscience) 470 nm excitation light was adjusted to approximately 80-100 µW at the tip of the patch cord (fiber core diameter: 400 µm; Doric Lenses). Fluorescence emission was passed through a 535 nm bandpass filter and focused onto the complementary metal-oxide semiconductor (CMOS) camera sensor through a tube lens. Samples were collected at 20 Hz, interleaved, sequentially between 470 and 415 nm excitation channels, using a custom Bonsai^69^ workflow. Time stamps of task events were collected simultaneously through an additional synchronized camera aimed at the Med Associates interface, which sent light pulses coincident with task events. Signals were saved using Bonsai software and exported to MATLAB (MathWorks, Natick, MA) for analysis. Recordings were collected unilaterally from the hemisphere with the strongest fluorescence signal at the start of the experiment.

### Optogenetic inhibition of VTA_DA_→BLA terminals at cue onset during Pavlovian-to-instrumental transfer and outcome-specific devaluation tests

#### Subjects

Thirty male (*N* = 14) and female (*N* = 16) transgenic Th-cre+ (hemizygous) Long Evans rats aged approximately 10 weeks at the time of surgery were used in this study to assess the necessity of cue-evoked VTA_DA_→BLA projection activity for the adaptive decision making during the Pavlovian-to-instrumental transfer and outcome devaluation tests. Seventeen (7 males) served in the experimental group and 13 (7 males) served as controls. Subjects with misplaced optic fibers or viral expression (*N* = 7) were excluded from the dataset.

#### Surgery

Th-cre rats were randomly assigned to a viral group and infused bilaterally with a cre-dependent AAV encoding either the inhibitory opsin archaerhodopsin T (ArchT; *N* = 17; 7 males; AAV5-CAG-FLEX-ArchT-tdTomato, Addgene) or a tdTomato fluorescent protein control (tdTomato; *N* = 11; 7 males; AAV5-CAG-FLEX-tdTomato, Addgene). Virus (0.4 µl) was infused bilaterally at a rate of 0.1 µl/min into the VTA (AP: −5.3; ML: ±0.75; DV: - 8.35 mm from bregma) using a 28-gauge injector. Injectors were left in place for 10 min following infusion. Optical fibers (200-µm core, 0.39 NA, Thorlabs, Newton, NJ) held in ceramic ferrules (Kientec Systems, Stuart, FL) were implanted bilaterally in the BLA (AP: −2.7; ML: ±5.0; DV: −8.6 mm from bregma). Experiments commenced 4-5 weeks after surgery to allow sufficient expression in VTA_DA_→BLA terminals at the time of manipulation (7-9 weeks post-surgery).

#### Optogenetic inhibition of VTA_DA_→BLA projections during Pavlovian-to-instrumental transfer and outcome-specific devaluation probe tests

Rats received magazine training, instrumental and Pavlovian conditioning, Pavlovian-to-instrumental transfer test, retraining, conditioned taste aversion, and the outcome-specific devaluation probe test as above. During the last 3 sessions of Pavlovian conditioning and instrumental retraining were attached to the optical tether (200 µm, 0.22 NA, Doric Lenses), but no light was delivered. Optogenetic inhibition was used to attenuate the activity of ArchT-expressing VTA_DA_ axons and terminals in the BLA at the time of each cue onset during the Pavlovian-to-instrumental transfer and outcome-specific devaluation probe tests. Green light (532 nm; 10 mW) was delivered to the BLA via a laser (Changchun New Industries Optoelectronics Technology Co.) connected through a ceramic mating sleeve (Thorlabs) to the ferrule implanted on the rat. During the PIT and devaluation probe tests, 2.5 s of continuous light was delivered beginning 0.5 s before each cue onset. Light effects were estimated to be restricted to the BLA core based on predicted irradiance values (https://web.stanford.edu/group/dlab/cgi-bin/graph/chart.php). Following PIT prior to devaluation 2 subjects were removed from the control group as a result of head cap detachment and post consumption test failure and, therefore, are not included in the devaluation dataset.

### Pharmacological inactivation of dopamine D1 receptors during Pavlovian-to-instrumental transfer test

#### Subjects

Twenty six male (*N* = 15) and female (*N* = 11) Long Evans rats aged approximately 10 weeks at the time of surgery were used in this study to assess the necessity of BLA dopamine D1 receptor activity for the adaptive decision making during the Pavlovian-to-instrumental transfer test. Nine (5 males) rats served in the vehicle control group, ten (6 male) served in the 0.5 µg SCH-23390 group, and seven (4 male) served in the 1.0 µg SCH-23390 group. Subjects with misplaced cannula (*N* = 2), or lesions in the infusion region (*N* = 4) were excluded from the dataset.

#### Surgery

For all rats, 22-gauge stainless-steel guide cannulae (Plastics One) were implanted bilaterally targeted 1 mm above the BLA (AP: −2.7; ML: ±5.0; DV: −7.6 mm from bregma). Behavioral training commenced ∼1 week after surgery to allow for recovery.

#### Drug administration

The dopamine D1 receptor antagonist, SCH-23390 (Sigma Aldrich, Burlington, MA), was dissolved in sterile saline vehicle to a concentration of 0.5 or 1 µg/0.5µl. Drug or vehicle was infused bilaterally into the BLA in a volume of 0.5 µl over 1 min via injectors inserted into the guide cannulae fabricated to protrude 1 mm ventral to the cannula tip by using a microinfusion pump. Injectors were left in place for at least one additional minute to ensure full infusion. This infusion volume was selected to avoid spread to the adjacent central nucleus of the amygdala^68, 70, 71^. Rats were placed in the conditioning chamber and testing began ∼10 min following infusion to allow sufficient time for the drug to become effective. Doses were selected based on evidence of their prior efficacy in appetitive tasks when infused into the BLA^72, 73^.

#### Pharmacological inactivation of dopamine D1 receptors during Pavlovian-to-instrumental transfer test

Rats received magazine training, instrumental and Pavlovian conditioning, Pavlovian-to-instrumental transfer test as above. SCH-23390 was infused into the BLA to temporarily inactivate dopamine D1 receptors during the Pavlovian-to-instrumental transfer test.

### Optogenetic inhibition of VTA_DA_→BLA terminals at cue onset during Pavlovian blocking with Pavlovian-to-instrumental transfer test

#### Subjects

Twenty-one male (*N* = 11) and female (*N* = 10) transgenic Th-cre+ (hemizygous) Long Evans rats (*N* = 18) and Th-cre- littermates (*N* = 3) aged approximately 8-10 weeks at the time of surgery were used in this study to assess the necessity of cue-evoked VTA_DA_→BLA projection activity for the ability of cue-reward predictions to constrain learning. Ten (5 males) served in the experimental group and 11 (6 males; 8 Th-cre+, 3 Th-cre) served as controls. Subjects with misplaced optic fibers or viral expression (*N* = 2) were excluded from the dataset.

#### Surgery

Th-cre+ rats were randomly assigned to a viral group and infused bilaterally with a cre-dependent AAV encoding either ArchT (*N* = 10; 5 males; AAV5-CAG-FLEX-ArchT-tdTomato, Addgene) or a tdTomato fluorescent protein control (tdTomato; *N* = 8; 4 males; AAV5-CAG-FLEX-tdTomato, Addgene). Th-cre- rats were assigned to the control group and infused bilaterally with a cre-dependent AAV encoding ArchT (*N* = 3; 2 males; AAV5-CAG-FLEX-ArchT-tdTomato, Addgene). Virus (0.4 µl) was infused bilaterally at a rate of 0.1 µl/min into the VTA (AP: −5.3; ML: ±0.75; DV: −8.35 mm from bregma) using a 28-gauge injector. Injectors were left in place for 10 min following infusion. Optical fibers (200-µm core, 0.39 NA, Thorlabs, Newton, NJ) held in ceramic ferrules (Kientec Systems, Stuart, FL) were implanted bilaterally in the BLA (AP: −2.7; ML: ±5.0; DV: −8.6 mm from bregma). Experiments commenced 4-5 weeks after surgery to allow sufficient expression in VTA_DA_→BLA terminals at the time of manipulation (7-9 weeks post-surgery).

#### Optogenetic inhibition of VTA_DA_→BLA projections during compound conditioning

Rats received magazine training, instrumental conditioning, and visual cue Pavlovian conditioning as above. During these conditioning sessions rats were attached to the optical tether (200 µm, 0.22 NA, Doric Lenses), but no light was delivered. Optogenetic inhibition was used to attenuate the activity of ArchT-expressing VTA_DA_ axons and terminals in the BLA at the time of each cue onset during compound conditioning. Green light (532 nm; 10 mW) was delivered to the BLA via a laser (Changchun New Industries Optoelectronics Technology Co.) connected through a ceramic mating sleeve (Thorlabs) to the ferrule implanted on the rat. During each compound conditioning sessions, 2.5 s of continuous light was delivered beginning 0.5 s before each cue onset. Following compound conditioning, rats proceeded to the PIT test as described above, during which they were tethered to the optical patch cords, but no light was delivered.

### Histology

Following behavioral experiments, rats were deeply anesthetized with Nembutal and transcardially perfused with phosphate buffered saline (PBS) followed by 4% paraformaldehyde (PFA). Brains were removed and post-fixed in 4% PFA overnight, placed into 30% sucrose solution, then sectioned into 50-μm slices using a cryostat and stored in cryoprotectant. Immunofluorescence was used to confirm expression of GRAB_DA_ in the BLA. Floating coronal sections were washed 3 times in 1x PBS for 30 min and then blocked for 1–1.5 hr at room temperature in a solution of 3% normal goat serum and 0.3% Triton X-100 dissolved in PBS. Sections were then washed 3 times in PBS for 15 min and incubated in blocking solution containing chicken anti-GFP polyclonal antibody (1:1000; Abcam, Cambridge, MA) with gentle agitation at 4°C for 18–22 hr. Sections were next rinsed 3 times in PBS for 30 min and incubated with goat anti-chicken IgY, Alexa Fluor 488 conjugate (1:500; Abcam) in blocking solution at room temperature for 2 hr. Sections were washed a final 2 times in PBS for 10 min. Slices were rinsed in a DAPI solution for 7 min (5 mg/mL stock, 1:10000), washed 3 times in PBS for 15 min, mounted on slides and coverslipped with ProLong Gold mounting medium.

To confirm the expression of GCamp6s in the VTA_DA_ neurons GFP fluorescence with a TH co-stain was used. Floating coronal sections were washed 3 times in 1x PBS for 30 min and then blocked for 1–1.5 hr at room temperature in a solution of 3% normal goat serum and 0.3% Triton X-100 dissolved in PBS. Sections were then washed 3 times in PBS for 15 min and incubated in blocking solution containing rabbit anti-TH polyclonal antibody (1:1000; Abcam, Cambridge, MA) with gentle agitation at 4°C for 18–22 hr. Sections were next rinsed 3 times in PBS for 30 min and incubated with goat anti-rabbit, Alexa Fluor 594 conjugate (1:1000; Abcam) in blocking solution at room temperature for 2 hr. Sections were washed a final 2 times in PBS for 10 min. Slices were rinsed in a DAPI solution for 7 min (5 mg/mL stock, 1:10000), washed 3 times in PBS for 15 min, mounted on slides and coverslipped with ProLong Gold mounting medium.

tdTomato fluorescence with a Th costain was used to confirm expression of ArchT-tdTomato in VTA_DA_ neurons. Floating coronal sections were washed 3 times in 1x PBS for 30 min and then blocked for 2 hr at room temperature in a solution of 3% normal donkey serum and 0.2% Triton X-100 dissolved in PBS. Sections were then washed 3 times in PBS for 15 min and incubated in blocking solution containing rabbit anti-TH antibody (1:1000; EMD Millipore, Burlington, MA) with gentle agitation at 4°C for 44-48 hr. Sections were next rinsed 3 times in PBS for 30 min and incubated with goat anti-rabbit IgG, Alexa Fluor 488 conjugate (1:500; Thermofisher Scientific, Waltham, MA) in blocking solution at room temperature for 2 hr. Sections were washed a final 2 times in PBS for 10 min. Immunofluorescence was also used to confirm expression of ArchT-tdTomato in axons and terminals in the BLA. Floating coronal sections were washed 2 times in 1x PBS for 10 min and then blocked for 2 hr at room temperature in a solution of 10% normal goat serum and 0.5% Triton X-100 dissolved in PBS. Sections were then washed 3 times in PBS for 15 min and incubated in blocking solution containing rabbit anti DsRed polyclonal antibody (1:1000; EMD Millipore, Burlington, MA) with gentle agitation at 4°C for 18-22 hr. Sections were next rinsed 3 times in blocking solution for 30 min and incubated with goat anti-rabbit IgG, Alexa Fluor 594 conjugate (1:500; Thermofisher Scientific) in blocking solution at room temperature for 2 hr. Sections were washed a final 2 times in PBS for 10 min.

Images were acquired using a Keyence BZ-X710 microscope (Keyence, El Segundo, CA) with a 4x, 10x, and 20x objective (CFI Plan Apo), CCD camera, and BZ-X Analyze software.

### Data analysis

#### Behavioral analysis

Behavioral data were processed with Microsoft Excel (Microsoft, Redmond, WA) and MATLAB (MathWorks). Press rates on the last 2 sessions of instrumental training were averaged across levers then across days and compared between groups to test for any pre-existing group differences in instrumental behavior. For Pavlovian conditioning, conditional food-port approach responses during the Pavlovian and compound conditioning sessions were assessed by comparing the rate of entries into the food-delivery port (entries/min) during the 30-s cue periods relative to the 30-s baseline periods prior to cue onset (preCue). Data were averaged across trials for each cue and then averaged across the two cues. We averaged data across the final 2 conditioning sessions. For PIT tests, entry rate into the food port during the 30-s cues was compared to the baseline 30-s preCue periods. Data were averaged across trials for each cue and then averaged across cues. Lever press rates (presses/min) during the 30-s baseline preCue periods were compared to that during the 30-s cue periods to capture the cue-induced change in lever pressing. During the cue, lever presses were separated for presses on the lever that, during training, earned the same outcome as the upcoming or presented cue (Same presses) versus those on the other available lever (Different presses). In all cases, Same and Different presses did not significantly during the pre-cue baseline period (lowest *P* = 0.26) and so were collapsed into a single baseline press rate. Data was averaged across trials, then averaged across cue types. To evaluate the selectivity of pressing towards the lever associated with the same outcome as predicted by the cue, we computed a selectivity index [Same presses/(Same + Different presses)] or score (Same presses – different presses). For the devaluation test, we evaluated entries into the food-delivery port for a 35-s cue window combining the 30-s cue and 5-s post-cue period. This allowed us to capture behavior during the cue itself and the post-cue period, when reward was delivered during training, and, thereby capture cue-evoked entries and sensitivity to devaluation of the predicted outcome that occurs in controls (see Extended Data Figure 2-1). Food-port entries did not differ between the periods before the valuedf devalued cue (lowest *P* = 0.51) and so were collapsed to form a single baseline, which was compared to entries during the valued and devalued cues. To evaluate sensitivity to devaluation we computed a value index [Valued CS entries/ (Valued + Devalued CS entries)].

#### Fiber photometry analysis

Data were pre-processed using a custom-written pipeline in MATLAB (MathWorks, Natick, MA)^16, 36^. The 415-nm and 470-nm signals were fit using an exponential curve. Change in fluorescence (ΔF/F) at each time point was calculated by subtracting the fitted 415-nm signal from the 470-nm signal and normalizing to the fitted 415-nm data [(470-fitted 415)/fitted 415)]. Non-normalized 470-nm signal is provided in Extended Data Figure 1-1. The ΔF/F data was corrected for signal decay using a second order polynomial function and Z-scored to the average of the whole session [(ΔF/F - mean ΔF/F)/std(ΔF/F)]. Z-scored traces were then aligned to behavioral event timestamps. Individual trial data were excluded if the patch cord became detached or if artifactual signal due to excessive motion or patch cord twisting was detected (n = 3 PIT trials from 3 subjects (1 trial/subject)).

Using a waveform analysis^74^, event-related activity was analyzed 60 s before to 60 s after cue onset. To ensure independence of observations, the analysis was performed at the subject-level average waveform. For the PIT test data, GrabDA2m and GFP cue-aligned signals, were independently averaged to generate time series for each subject. For the devaluation test data, for each subject, the Valued cue and Devalued cue trials were independently averaged to generate a mean time-series for each CS type for each subject. The observed difference waveform (mean GrabDA2m cue – mean GFP cue or mean Valued cue – mean Devalued cue) was analyzed for statistical significance at every time point using two non-parametric approaches: a two-sample permutation test and a bootstrap confidence interval. For the permutation test, with 1000 iterations a null distribution of condition differences was generated by randomly assigning the cue condition labels and calculating the difference of the resampled means across the entire time window. The *P* value at each time point was the proportion of permutations whose mean difference values were more extreme than the observed difference between the actual samples. A time point was flagged as significant if *P* < 0.05. To provide a robust estimate of the true population mean difference, a matrix of 1000 bootstrapped means were generated by resampling with replacement the subject-level average waveform for both GrabDA2m and GFP or Valued cue and Devalued cue. The difference between cues was calculated for each of the 1000 iterations. CI for each timepoint were percentiles at that timepoint of the bootstrap matrix (95%: 2.5, 97.5 percentiles), which were then expanded by a factor of √n/(n−1) to counter small sample narrowness bias. A significant difference was flagged whenever the CI did not contain the null of 0. For both approaches, a threshold of 10 consecutive time points was applied to the initial set of significant indices to correct for isolated, random fluctuations. Using this approach, we found significance increases in cue-evoked dopamine and differences between the Value and Devalued cues immediately after CS onset. Thus, to further quantify dopamine fluctuations, we calculated the Area under the curve (AUC) in the 4 s period immediately following cue onset and compared to the 4 s pre-cue baseline periods using a trapezoidal function. Quantifications and signal aligned to events were averaged across trials within a session and compared between groups.

Because lever pressing and/or entries into the food-delivery port can freely occur during cue presentation, we used a linear regression model to parse the extent to which dopamine fluctuations were explained by these discrete, but overlapping events ^49, 50^. To calculate kernels that correspond to the isolated response to each behavioral event for each cue type Cue onset (for devalued separating value vs. devalued), Cue offset, lever presses, and food-port entries, the Z-scored GRAB_DA_ signal 2 s prior to and 2 s after cue onset was modeled as the sum of the response to each event. Discrete events cue onset, offset, lever press, and food-port entry were convolved into a time series representing the time of the event in a series of values between 0 – 1 using a 4-s gaussian distribution. For cue onset and offset, we used a half-gaussian convolution (0 s – 4 s). Because behavioral events include both movements to make the action and a following outcome (e.g., non-reinforcement), the press and food-port entry convolution was centered on the event (−2 – 2 s). For each cue type for each subject, the coefficients of the kernels were solved using the method of least-squares with the MATLAB function *regress*. The kernel output for the 2 s prior to and 2 s after cue onset was averaged across trials for each cue type for each subject. To calculate the percentage of variance explained by each regressor class, the predicted dopamine signal was recalculated removing one regressor class (e.g., cue) at a time. The percent variance explained by the removed regressor was calculated by subtracting the variance explained by the model with the regressor removed from the explained variance of the full model, and expressing this as a percentage of the variance explained by the full model [(*v*_full_ - *v*_partial_/*v*_full_)*100).

#### Statistical analysis

Datasets were analyzed by two-tailed, paired and unpaired Student’s *t* tests, two-, or three-way repeated-measures analysis of variance (ANOVA), and simple linear regression as appropriate (GraphPad Prism, GraphPad, San Diego, CA; SPSS, IBM, Chicago, IL). For the few datasets that were slightly non-normal, results were cross-checked using non-parametric statistics and the findings were identical. We opted to use parametric statistics for consistency across experiments and given evidence that ANOVA is robust to slight non-normality^75,76^. All other *post hoc* tests were corrected for multiple comparisons using the Bonferroni method and used to clarify main and interaction effects. Alpha levels were set at *P* < 0.05.

### Sex as a biological variable

Male and female rats were used in approximately equal numbers for each experiment, but the *N* per sex was underpowered to examine sex differences. Sex was therefore not included as a factor in statistical analyses, though individual data points are visually disaggregated by sex.

### Rigor and reproducibility

Group sizes were estimated *a priori* based on prior work using male Long Evans rats in this behavioral task^16, 38, 62, 77^ and to ensure counterbalancing of Cue-reward and Lever-outcome pairings. Investigators were not blinded to viral group because they were required to administer virus. All behaviors were scored using automated software (MedPC). Each experiment included at least 1 replication cohort and cohorts were balanced by viral group, Cue-reward and Lever-reward pairings, hemisphere etc. prior to the start of the experiment.

## Data availability

All data that support the findings of this study is available as supplemental files.

## Code availability

Custom-written MATLAB code is available from the corresponding author upon request and the basic code is available via Dryad (https://doi.org/10.5068/D1109S).

## ACKNOWLEDGEMENTS

This research was supported by NIH grants DA057084 (KMW & MJS) and DA054967 (MJS), the Wendell Jeffrey and Bernice Wenzel Term Chair in Behavioral Neuroscience to KMW, the Staglin Center for Behavior and Brain Sciences, and NHMRC Investigator Grant 2024/GNT2034225 (MJS).

## AUTHOR CONTRIBUTIONS

KMW and KR designed the research, analyzed, and interpreted the data. KR conducted the research with assistance from CF, AM, JA, JQ, NKG. AS contributed to the standardization of the MATLAB analysis pipeline. MJS contributed to the conception and design and advised on manuscript. KMW and KR wrote the manuscript.

## EXTENDED DATA

**Extended data figure 1-1.**
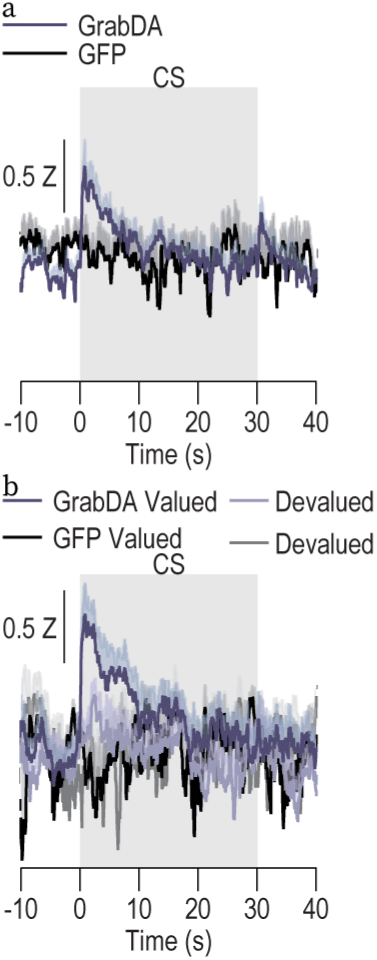
Cue-evoked BLA dopamine without 415 nm signal subtraction. GRAB_DA_ or control GFP fluorescence changes (drift-corrected 470 nm without subtraction of 415 nm signal) in response to cue presentation during the PIT (a) or devaluation (b) test. Data presented as mean ± s.e.m. As shown in the main data, the onset of both cues triggered robust, transient dopamine release that was larger for cue signaling the valued reward than the cue signaling the devalued reward. PIT, GRABDA: N = 10 (7 male); GFP: N = 4 (2 male). Devaluation, GRABDA: N = 10 (7 male); GFP: N = 4 (2 male).

**Extended data figure 1-2.**
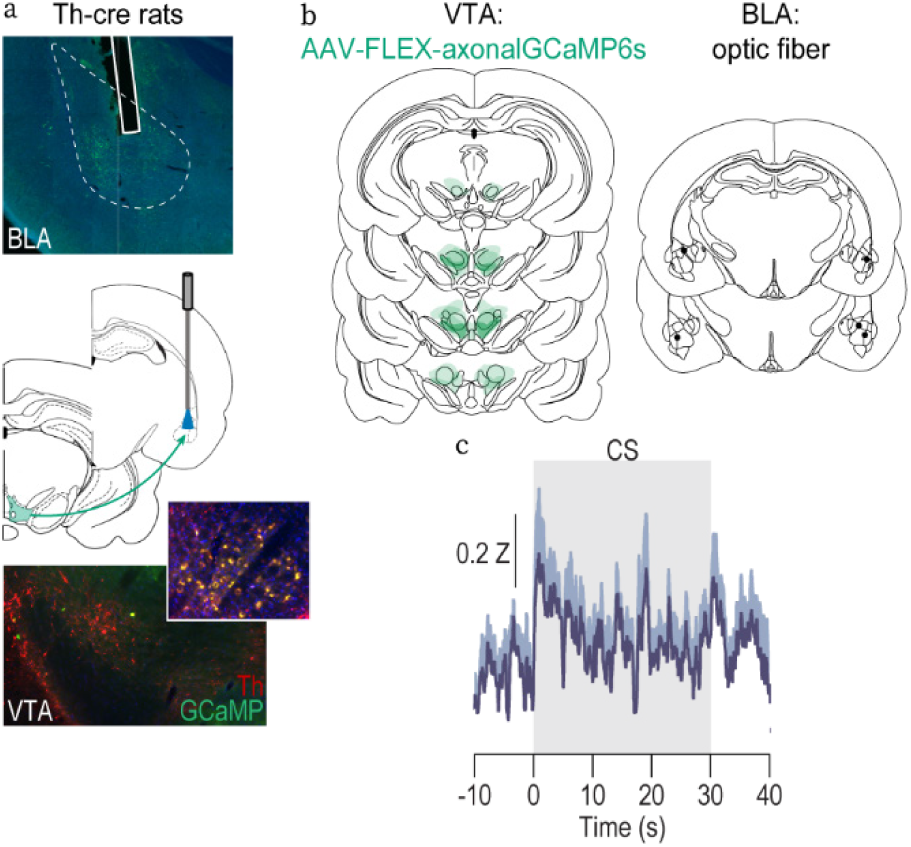
Cue-evoked activity of VTA_DA_ axons and terminals in BLA. Male and female rats were food restricted and received instrumental conditioning in which each of two different lever-press actions earned one of two distinct food rewards (e.g., left press→ensure/right press→pellets; 11 sessions) and then received Pavlovian conditioning during which 2 distinct, 30-s auditory cues (aka, conditioned stimuli) each predicted delivery of one of the unique food rewards at cue offset (e.g., white noise—ensure/click—pellets). We used fiber photometry to record fluorescent activity of the axonally-targeted calcium indicator GCaMP6s expressed in VTA_DA_ neurons during the PIT test. **(a)** Bottom: axonal-GCaMP6s expressed in VTA_DA_ neurons. Middle: Fiber photometry approach for imaging axonal-GCaMP6s fluorescence changes in VTA_DA_ axons and terminals in BLA. Top: Representative fluorescent image of BLA axonal-GCaMP6s expression and fiber placement. **(b)** Schematic representation of axonal-GCaMP6s expression in VTA and optical fiber tips in BLA for all subjects. **(c)** Axonal-GCaMP6s fluorescence changes (Z-score) during the cues the PIT test. *N* = 5 (3 male). Data presented as trial-averaged, between-subject mean ± s.e.m. Cue onset and offset triggered a transient increase in VTA_DA_ axonal activity release.

**Extended data figure 2-1.**
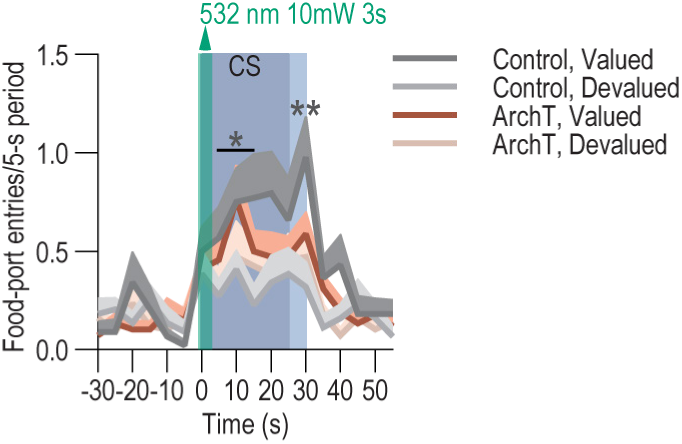
The effect of devaluation and VTA_DA_→BLA inhibition persists into the 5-s post-cue period. Food-port entries in 5-s bins before, during, and after cue (conditioned stimulus, CS) presentation. Blue, 30-s CS; Light blue, 5-s post-CS period. 3-way repeated measures ANOVA, Time x Virus x Value: F_17,442_ = 1.63, *P* = 0.05; Time: F_17,442_ = 14.71, *P* < 0.0001; Virus: F_1,26_ = 0.38, *P*=0.54; Value: F_1,26_ = 11.80, *P* = 0.002; Time x Virus: F_17,442_ = 0.52, *P* = 0.94; Time x Value: F_17,442_ = 3.33, *P* < 0.0001; Virus x Value: F_1,26_ = 3.61, *P* = 0.07. The devaluation effect in controls persisted into the 5-s period after CS offset during which, in training, the reward was delivered and consumed. Devaluation: Control *N* = 11 (7 male); ArchT *N* = 17 (7 male). \**P* < 0.05, \*\**P* < 0.01 Control Valued vs. Devalued, Bonferroni corrected.

**Extended Data Figure 2-2.**
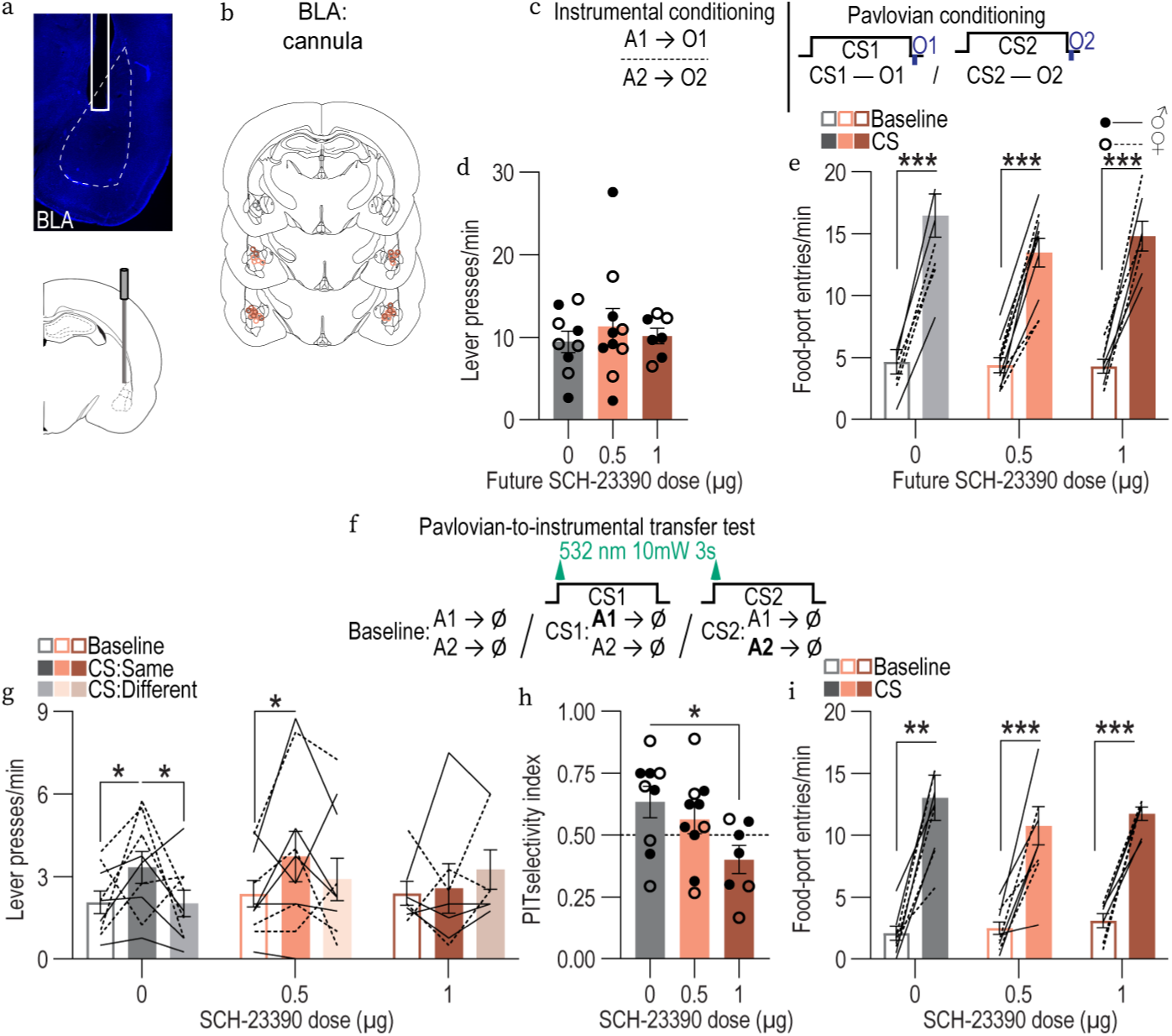
BLA dopamine D1 receptor activation mediates the influence of cue-reward predictions on decision making. If BLA dopamine supports cue-reward predictions, we reasoned that inhibiting BLA dopamine receptors should, like optogenetically inhibiting cue-evoked VTA_DA_→BLA activity, prevent reward-predictive cues from guiding decision making. We focused on BLA D1 receptors because D1-family receptors are commonly viewed as well suited to detecting the larger dopamine elevations produced by phasic dopamine release^78^ like that triggered by reward-predictive cues. Rats first received instrumental conditioning, without manipulation, in which each of two different lever-press actions earned one of two distinct food rewards. Rats then received Pavlovian conditioning, also without manipulation, during which 2 distinct 30-s auditory cues each predicted delivery of one the unique food rewards at cue offset. They were then given a PIT test during which the levers were available and each cue was presented 4 times in pseudorandom order to assess its influence over action selection. Ten minutes prior to this test we infused the dopamine D1 receptor antagonist, SCH-23390, into the BLA. **(a)** Bottom: Guide cannula were implanted bilaterally above BLA. Top: Representative image of guide cannula targeting BLA. **(b)** Schematic representation of injector tip placements in BLA for all subjects. **(c-e)** Instrumental and Pavlovian conditioning **(c)** Conditioning procedures. A, action (left or right lever press); CS, 30-s conditioned stimulus (aka, “cue”, white noise or click) followed immediately by reward outcome (O, ensure solution or grain pellet). **(d)** Lever-press rate averaged across levers and across the final 2 instrumental sessions. 1-way ANOVA, F_2, 23_ = 0.32, *P* = 0.73. **(e)** Food-port entry rate during averaged across cues and across the final 2 Pavlovian conditioning sessions. 2-way repeated measures ANOVA, CS: F_1, 23_ = 293.2, *P* < 0.0001; Drug group: F_2, 23_ = 0.72, *P* = 0.50; CS x Drug: F_2, 23_ = 1.86, *P* = 0.18. There were no pre-existing differences in either instrumental press rate or Pavlovian conditional goal approach responses between the future drug infusion groups. **(f-i)** Outcome-specific Pavlovian-to-instrumental transfer test. **(f)** PIT procedure. Ø, no reward was delivered. **(g)** Lever-press rates on the lever that earned the “Same” outcome as predicted or current cue or on the other available lever (Different) compared to pre-cue baseline press rate averaged across levers. Planned comparison 2-tailed students t-tests: Control: Baseline vs. Same, t_9_ = 1.96, *P* = 0.05, 95% CI - 0.03 – 2.56; Same v Different, t_9_ = 2.03, *P* = 0.048, 95% CI 0.009 – 2.60; 0.5µg: Baseline vs. Same, t_10_ = 2.21, *P* = 0.03, 95% CI 0.12 – 2.58; Same v Different, t_10_ = 1.35, *P* = 0.18, 95% CI −0.419 – 2.06; 1µg: Baseline vs. Same, t_7_ = 0.24, *P* = 0.81, 95% CI −1.29 – 1.65; Same v Different, t_7_ = 0.93, *P* = 0.36, 95% CI −2.15 – 0.79. **(h)** PIT selectivity index [Same presses/(Same + Different presses)]. 1-way ANOVA, F_2, 23_ = 3.48, *P* = 0.048. Inactivation of BLA dopamine D1 receptors attenuated the ability of the cues to bias action selection towards the lever associated with the same predicted reward than that associated with the different reward. **(i)** Food-port entry rate. 2-way repeated measures ANOVA, CS: F_1, 23_ = 120.2, *P* < 0.0001; Drug: F_2, 23_ = 0.34, *P* = 0.72; CS x Drug: F_2, 23_ = 1.04, *P* = 0.137. BLA dopamine D1 receptor inactivation did not disrupt expression of the conditional approach responses to the shared goal location, a behavior that does not require prediction of the identity of the specific reward. Control *N* = 9 (5 male); 0.5µg *N* = 10 (6 male); 1µg *N* = 7 (4 male). Data presented as trial-averaged, between-subject mean ± s.e.m. with individual data points. *P<0.05, **P<0.01, ***P<0.001.

